# psichomics: graphical application for alternative splicing quantification and analysis

**DOI:** 10.1101/261180

**Authors:** Nuno Saraiva-Agostinho, Nuno L. Barbosa-Morais

**Affiliations:** Instituto de Medicina Molecular João Lobo Antunes, Faculdade de Medicina, Universidade de Lisboa, Av. Professor Egas Moniz, 1649-028 Lisboa, Portugal

## Abstract

Alternative pre-mRNA splicing generates functionally distinct transcripts from the same gene and is involved in the control of multiple cellular processes, with its dysregulation being associated with a variety of pathologies. The advent of next-generation sequencing has enabled global studies of alternative splicing in different physiological and disease contexts. However, current bioinformatics tools for alternative splicing analysis from RNA-seq data are not user-friendly, disregard available exon-exon junction quantification or have limited downstream analysis features. To overcome such limitations, we have developed *psichomics*, an R package with an intuitive graphical interface for alternative splicing quantification and downstream dimensionality reduction, differential splicing and gene expression and survival analyses based on The Cancer Genome Atlas, the Genotype-Tissue Expression project and user-provided data. These integrative analyses can also incorporate clinical and molecular sample-associated features. We successfully used *psichomics* to reveal alternative splicing signatures specific to stage I breast cancer and associated novel putative prognostic factors.

## INTRODUCTION

Alternative splicing fosters transcriptome diversity in eukaryotes through the processing of pre-mRNAs from the same gene into distinct transcripts that may encode for proteins with different functions (1, 2). Alternative splicing is involved in multiple cellular processes, such as apoptosis and autophagy regulation (1, 2), and is especially prevalent in humans, where approximately 93% of genes display alternatively spliced transcripts whose regulation may differ across tissues and developmental stages (2–4). Consistently, alternative splicing dysregulation has been linked with cancer, neurodegeneration and other diseases (2, 5, 6). For instance, splicing alterations mediated by the key regulator SRSF1 may impact multiple hallmarks of cancer, such as resistance to apoptosis and tissue invasion (5).

The relevance of alternative splicing changes in physiological and disease conditions, along with the increasing economic feasibility of next-generation RNA sequencing (RNA-seq), has progressively driven transcriptome-wide alternative splicing studies (3, 7-10) and promoted large consortium efforts to assemble publicly accessible splicing data. Such consortia include The Cancer Genome Atlas (TCGA), that catalogues clinical and molecular profiling data from multiple human tumours (11), and the Genotype-Tissue Expression (GTEx) project, that focuses on profiling normal human multi-tissue data (12). Among the openly available processed data from these projects, counts of RNA-seq reads aligned to exon-exon junctions may be exploited for alternative splicing quantification and further analysis. Indeed, the ability to couple proper differential splicing analysis with, for instance, gene expression, protein domain annotation, clinical information or literature-based evidence enables researchers to extract, from those comprehensive public datasets, valuable insights into the role of alternative splicing in physiological and pathological contexts, as well as putative splicing-associated prognostic factors and therapeutic targets (7-10, 13).

Several tools are currently available to quantify, analyse and visualise alternative splicing data, including AltAnalyze (14), MISO (15), SpliceSeq (16), FineSplice (17), spliceR (18), VAST-TOOLS (19), rMATS (20), SUPPA (21), jSplice (22), JunctionSeq (23) and Whippet (BioRxiv: https://doi.org/10.1101/158519). However, each of such tools suffers from at least one of the following shortcomings:

– Lack of support for imputing pre-processed data, leading to redundant, time-consuming RNA-seq read alignment and exon-exon junction detection, preceding alternative splicing quantification when exon-exon junction quantification is already available (e.g. when analysing TCGA or GTEx data).
– Limited set of statistical options for differential splicing analysis, mostly relying on median-based non-parametric tests and restricted to pairwise comparisons.
– No incorporation of molecular or clinical information enabling analyses that reflect factorial designs or test linear models, for example. This is particularly limiting in the exploration of clinical datasets where, for instance, survival analyses permit assessing the potential prognostic value of alternative splicing events.
– No support for transcriptome-wide filtering and sub-setting of events, based on common features or the outcome of statistical analyses, for interactive exploration of individual events of interest.
– No user-friendly interactive graphical interface neither support for customisable statistical plots.

Moreover, to our knowledge, none of those tools currently incorporates support for survival analysis, exploratory and differential analyses of gene expression, or tests for association between gene expression levels and/or alternative splicing quantifications changes.

To offer a comprehensive pipeline that integrates all the aforementioned features through both a command-line and an easy-to-use graphical interface, we have developed *psichomics*, an R package to quantify, analyse and visualise alternative splicing and gene expression data using TCGA, GTEx and/or user-provided data. Our tool interactively performs dimensionality reduction, differential splicing and gene expression and survival analyses with direct incorporation of molecular and clinical features. We successfully employed *psichomics* to analyse stage I breast cancer TCGA data and identified alternative splicing events with putative prognostic value. *psichomics* is freely available in Bioconductor at http://bioconductor.org/packages/psichomics.

## MATERIALS AND METHODS

*psichomics* was developed as an R package with a modular design, allowing to easily modify and extend its components. These include support for multiple file formats and automatic data retrieval from external sources (e.g. TCGA and GTEx), parsing and standardisation of alternative splicing event identifiers from different programs and annotations and the implementation of a variety of data analysis methodologies.

The program’s workflow for alternative splicing analysis begins with the loading of splice junction read count data from the user’s computer or external sources, followed by the quantification of alternative splicing (in case no pre-computed quantification is loaded) and subsequent analyses. Alternative splicing quantification is based on RNA-seq reads that align to splice junctions and the genomic coordinates (annotation) of alternative splicing events. The proportion of reads aligned to junctions that support the inclusion isoform, known as the Percent Spliced-In or PSI (3), was the chosen quantification metric.

### Exon-exon junction quantification, gene expression and sample-associated data retrieval

Exon-exon junction and gene expression quantifications (obtained from pre-processed RNA-seq data) and clinical data are accessible through FireBrowse’s web application program interface (API) for TCGA data retrieval (http://firebrowse.org/api-docs). The FireBrowse API is used in *psichomics* to automatically download TCGA data according to the user-selected tumour type(s) as tab-delimited files within compressed folders, whose contents are subsequently loaded with minimal user interaction.

Contrastingly, GTEx does not currently provide any public API for automatic data retrieval, thus requiring the user to manually download exon-exon junction quantification, gene expression and clinical data from the GTEx website (http://gtexportal.org), for instance.

User-owned files may also be loaded in appropriate formats, as instructed when using *psichomics*, allowing for subsequent alternative splicing analysis from customised data.

### Gene expression pre-processing

Gene expression quantifications can be filtered based on user-provided parameters (for instance, to account solely for genes supported by 10 or more reads in 10 or more samples, as performed by default) and normalised by raw library size scaling using function *calcNormFactors* from R package *edgeR* (24). Afterwards, counts per million reads (CPM) are computed and log2-transformed (if desired) using the function *cpm* from *edgeR*. Log2-transformation is performed by default.

### Alternative splicing annotation

Annotations of alternative splicing events are available on-demand in *psichomics* for the Human hg19 (default) and hg38 genome assemblies. Custom annotation files can also be created by following the appropriate tutorial available at the package’s landing page in Bioconductor.

The hg19 annotation of human alternative splicing events was based on files used as input by MISO (15), VAST-TOOLS (19), rMATS (20), and SUPPA (21). Annotation files from MISO and VAST-TOOLS are provided in their respective websites, whereas rMATS and SUPPA identify alternative splicing events and generate such annotation files based on a given isoform-centred transcript annotation. As such, the human transcript annotation was retrieved from the UCSC Table Browser (25) in GTF and TXT formats, so that gene identifiers in the GTF file (misleadingly identical to transcript identifiers) were replaced with proper ones from the TXT version.

The collected hg19 annotation files were non-redundantly merged according to the genomic coordinates and orientation of each alternative splicing event and contain the following event types: skipped exon (SE), mutually exclusive exons (MXE), alternative first exon (AFE), alternative last exon (ALE), alternative 5´ splice site (A5SS), alternative 3´ splice site (A3SS), alternative 5´ UTR length (A5UTR), alternative 3´ UTR length (A3UTR), and intron retention (IR). The resulting hg19 annotation is available as an R annotation package in Bioconductor at http://bioconductor.org/packages/alternativeSplicingEvents.hg19, whereas the hg38 annotation (whose coordinates were converted from those of the hg19 annotation through function *liftOver* from package *rtracklayer* (26), based on the hg19 to hg38 chain file from UCSC) is also available as an R annotation package in Bioconductor at http://bioconductor.org/packages/alternativeSplicingEvents.hg38.

### Alternative splicing quantification

For each alternative splicing event in a given sample, its PSI value is estimated by the proportion of exon-exon junction read counts supporting the inclusion isoform therein (3). The junction reads required for alternative splicing quantification depend on the type of event (Figure 1). Alternative splicing events involving a sum of junction read counts supporting inclusion and exclusion of the alternative sequence below a user-defined threshold (10 by default) are discarded to avoid imprecise quantifications based on insufficient evidence.

**Figure 1.**
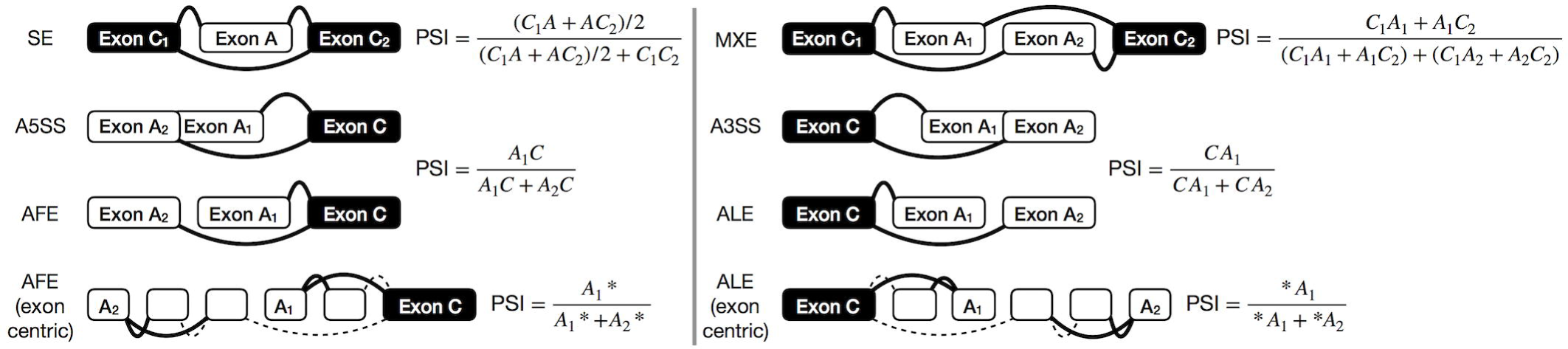
Splice junctions required to quantify alternative splicing based on event type. C_1_A and AC_2_ represent read counts supporting junctions between a constitutive (C_1_ or C_2_, respectively) and an alternative (A) exon and therefore alternative exon A inclusion, while C_1_C_2_ represents read counts supporting the junction between the two constitutive exons and therefore alternative exon A exclusion. A_1_* and A_2_* represent the sum of read counts supporting junctions spanning the alternative first (A_1_) and second (A_2_) exon, respectively. Legend: skipped exon (SE), mutually exclusive exons (MXE), alternative 5´ splice site (A5SS), alternative 3´ splice site (A3SS), alternative first exon (AFE) and alternative last exon (ALE).

Alternative splicing quantification in *psichomics* is currently based on exon-exon junction read counts, yet intron retention events require intron-exon junction read counts for their quantification (27), whereas alternative 5′-and 3′-UTR require exon body read counts. *psichomics* does not currently quantify those types of alternative splicing events.

By default, *psichomics* quantifies all skipped exon events. However, the user can select to measure other types of alternative splicing events (Figure 1) and may hand in the list of genes whose alternative splicing events are to be specifically quantified. Furthermore, the step of alternative splicing quantification may be avoided if previously performed. *psichomics* allows the user to save the quantification of alternative splicing in a file to be loaded in a future session.

### Data grouping

*psichomics* allows to group subjects and their samples or genes and their alternative splicing events for subsequent analysis. Subject and sample grouping can be performed based on available phenotypic (e.g. tissue type and histology) and clinical (e.g. disease stage, smoking history and ethnicity) features. Gene and splicing event grouping relies on respective user-provided identifiers. Moreover, the association between subject/sample groups specified by the user and those defined by the outcome of gene expression and alternative splicing analyses or by other clinical categorical variables can be statistically tested with Fisher’s exact tests, implemented through function *fisher.test* from *stats* (version 3.4.1).

### Dimensionality reduction

Dimensionality reduction techniques can be performed on alternative splicing and gene expression quantifications after centring and/or scaling the respective distributions (by default, they are only centred).

Principal component analysis (PCA) identifies the combinations of variables that contribute the most to data variance (28) and it is implemented through the singular value decomposition (SVD) algorithm provided by the *prcomp* function from R package *stats* (version 3.4.1). The total contribution of each variable (splicing event or gene) towards data variance along selected principal components is measured based on the implementation of *fviz_contrib* from *factoextra* (version 1.0.5).

Independent component analysis (ICA), a method used for decomposing data into statistically independent components (29), can also be performed through the *fastICA* function from the eponymous R package (version 1.2-1), preceded by data centring and/or scaling with the *scale* function.

As many of the aforementioned functions cannot handle missing data, a user-defined threshold for the accepted number of missing values per alternative splicing event or gene (5%, by default) is used to discard variables before performing dimensionality reduction, whereas the remaining missing values are imputed for each variable as the median from non-missing data samples.

Moreover, samples can be clustered using k-means, partitioning around medoids (PAM) or clustering large applications (CLARA) methods, with the latter being optimised for large datasets and thus preferred by default. The implementation of these methods is based on the *kmeans* function from *stats* (version 3.4.1) and *pam* and *clara* functions from *cluster* (version 2.0.6), respectively.

### Survival analysis

Kaplan-Meier estimators (and illustrating curves) (30) and proportional hazard (PH) models (31) may be applied to groups of patients defined by the user based on clinical features derived, for instance, from TCGA and user-owned data (GTEx features no survival data), with survival distributions being compared using the log-rank test. Survival analyses are implemented in *psichomics* using functions *Surv, survfit, survdiff* and *coxph* from R package *survival* (32).

To evaluate the prognostic value of a given alternative splicing event, survival analysis can be performed on groups of patients separated based on a given alternative splicing quantification (i.e. PSI) cut-off. Patients with multiple samples are assigned the average PSI value of their respective samples after sample filtering (e.g. when using TCGA data, only tumour samples are used for survival analysis by default). *psichomics* suggests an optimal cut-off that minimises the p-value of the log-rank test used to compare survival distributions. That optimisation employs the Brent method available in the *optim* function from the *stats* package (version 3.4.1). Survival analysis can also be performed on groups defined by an expression cut-off for a selected gene.

### Differential splicing and gene expression analyses

In *psichomics*, analysis of differential splicing between user-defined groups of samples can be performed on all or selected alternative splicing events. Given the non-normal distribution of PSI values (33, 34), median- and variance-based non-parametric tests, such as the Wilcoxon rank-sum (also known as Mann-Whitney U), Kruskal-Wallis rank-sum and Fligner-Killeen tests, are available and recommended (35). Levene’s and unpaired t-tests can nonetheless be performed as well. All these tests are available through the *stats* package (version 3.4.1) with their default settings, except for Levene’s test that was implemented based on the *leveneTest.default* function from the *car* package (version 2.1-6).

To correct for multiple testing where applicable, p-value adjustment methods for the family-wise error rate (Bonferroni, Holm, Hochberg and Hommel corrections) and the false discovery rate (Benjamini-Hochberg and Benjamini-Yekutieli methods) are available through function *p.adjust* from package *stats* (version 3.4.1). By default, multiple testing correction is performed using the Benjamini-Hochberg method.

Although the aforementioned statistical tests are also available to analyse the expression of single genes, genome-wide differential gene expression analysis is implemented based on gene-wise linear model fitting (using *lmFit* from R package *limma* (36)) for two selected groups, followed by moderated t-tests and the calculation of log-odds of differential expression, using empirical Bayes moderation of standard errors (function *eBayes* from *limma)* and gene-wise variance modelling *(limma-trend)*.

Statistical results can be subsequently explored through density and volcano plots with customisable axes to assist in the identification of the most significant changes when analysing distributions across single or multiple events, respectively. A corresponding table with the results of all statistical analyses is also available and can be retrieved as a tab-delimited plain text file.

### Correlation between gene expression and alternative splicing quantifications

The Pearson product-moment correlation coefficient, Spearman’s *rho* (default) and Kendall’s *tau*, all available with *cor.test* from *stats* (version 3.4.1), can be used to correlate gene expression levels with alternative splicing quantifications. Such analyses allow, for instance, to test the association between the expression levels of RNA-binding proteins (RBPs) and PSI levels of interesting splicing events to identify which of these may undergo RBP-mediated regulation. As such, a list of RBPs is provided in-app (37), but the user can also define their own group of genes of interest for the test.

### Gene, transcript and protein annotation and literature support

The representational state transfer (REST) web services provided by Ensembl (38), UniProt (39), the Proteins API (40) and PubMed (41) are used in order to annotate genes of interest with relevant biomolecular information (e.g. genomic location, associated transcript isoforms and protein domains, etc.) and related research articles. *psichomics* also provides the direct link to the cognate entries of relevant external databases, namely Ensembl (42), GeneCards (43), the Human Protein Atlas (44), the UCSC Genome Browser (45), UniProt (39) and VAST-DB (46).

### Performance benchmarking

To measure the time taken by *psichomics* to load data, normalise gene expression, quantify PSIs for skipped exon events and perform global differential expression and splicing analyses between pairs of GTEx tissues and between normal and primary solid tumour samples from multiple TCGA cohorts, the program was run 10 times with the same settings for different combinations of normal human tissues and tumour types in a machine running OS X 10.13.1 with 4 cores and 8GB of RAM, using Safari 11.0.1, RStudio Desktop 1.1.383 and R 3.4.1. The median duration of the 10 runs was used as the performance indicator.

To determine the approximate time complexity of the aforementioned steps in *psichomics*, gene expression and exon-exon junction quantification datasets were prepared based on approximate distributions obtained from the respective TCGA datasets: negative binomial distributions with a dispersion parameter of 0.25 and 0.2 reads and a mean parameter of 2000 and 100 reads for raw gene expression and exon-exon junction quantification, respectively. Each run was performed on datasets with numbers of samples ranging from 100 to 2500 in intervals of 100 (i.e. 100, 200, 300, …, 2500) and 20000 genes or 200000 splice junctions (gene expression or exon-exon junction quantification, respectively). Splice junction identifiers (required for alternative splicing quantification) were randomly retrieved from the TCGA reference annotation. Based on their respective read counts, around 9000 alternative splicing events (i.e. those for which all involved inclusion and exclusion junctions were retrieved) were quantified across selected samples per run. For differential gene expression and splicing analyses, samples were randomly divided into two groups based on the emitted values of a Bernoulli distribution with a probability of success of 50%.

Polynomials of orders 1 to 6 were fitted to the relation between running time and the number of samples. As the running time is assumed to always increase with an increasing number of analysed samples, fitted polynomials were constrained to be monotone for 0 or more samples, using function *monpol* from R package *MonoPoly* (47). The best polynomial fits (Figure 3) were selected based on analyses of variance (ANOVA) between fitted polynomials of consecutive orders, starting with the comparison between polynomials of orders 1 and 2. A polynomial with higher order is only selected if exhibiting a significantly better fit (p-value < 0.05).

**Figure 3.**
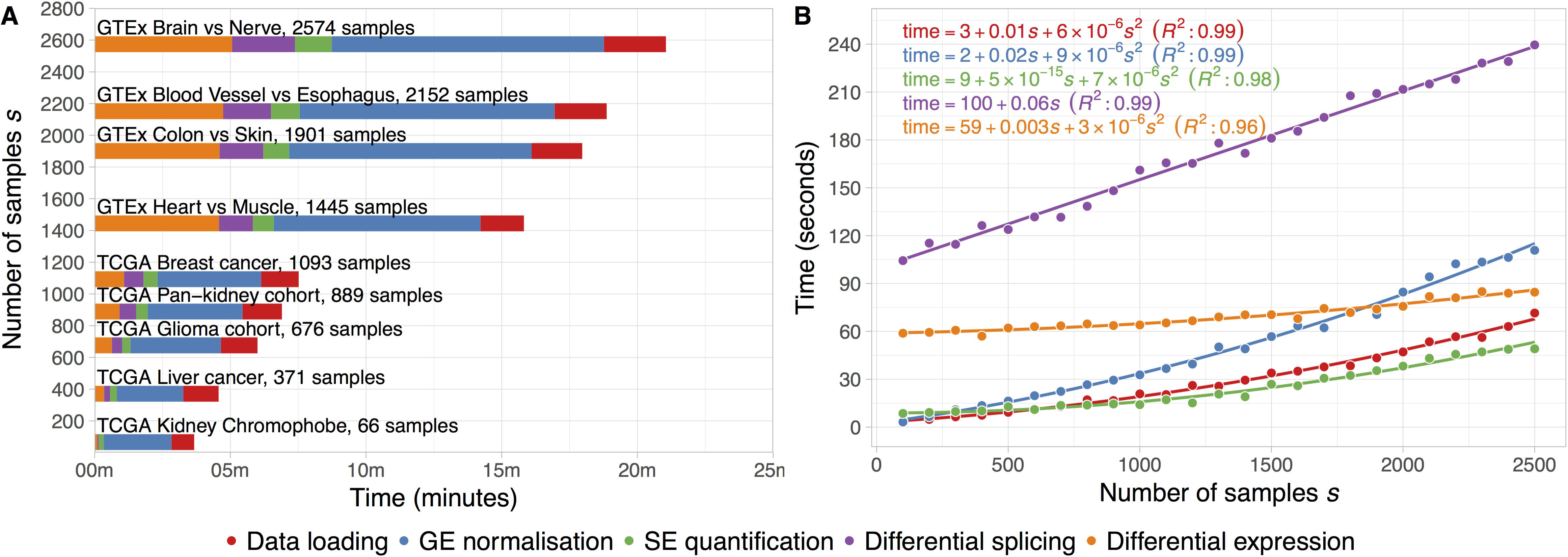
Performance benchmark for alternative splicing analysis using RNA-seq data from multiple TCGA and GTEx sample types. **A -** Median times of 10 runs of data loading, gene expression (GE) normalisation, skipped exon (SE) event quantification and differential expression and splicing analysis (normal versus tumour for TCGA data or pairwise tissue comparison for GTEx data) using *psichomics*. The default settings were used during the runs. **B -** Estimation of the time complexity of each of the aforementioned steps in *psichomics*. Randomly generated synthetic datasets of different sample size s were used as input. Equations and coefficient of determination (R^2^) for the best fits are displayed.

## RESULTS

*psichomics* offers both a graphical and a command-line interface. Although most features are common to both interfaces, we recommend less experienced users to opt for the graphical interface based on the *shiny* package (version 1.0.5), a web application framework available for R. To start the graphical interface, the user is required to load the *psichomics* package in R and run function *psichomics()*, resulting in the automatic launch of the user’s default web browser and of the program’s graphical interface as a local web app.

### Case study: exploration of clinically-relevant, differentially spliced events in breast cancer

Breast cancer is the cancer type with the highest incidence and mortality in women (48) and multiple studies have suggested that transcriptome-wide analyses of alternative splicing changes in breast tumours are able to uncover tumour-specific biomarkers (7, 8, 13). Given the relevance of early detection of breast cancer to patient survival, we used *psichomics* to identify novel tumour stage-I-specific molecular signatures based on differentially spliced events.

For the purposes of this case study, default *psichomics* settings were used unless otherwise stated. The analysis steps summarised below are easily reproducible by following the tutorials in the Bioconductor landing page for *psichomics*.

Alternative splicing quantification of the most recent TCGA breast cancer processed RNA-seq data available (2016-01-28) was performed by *psichomics* for skipped exons, mutually exclusive exons, alternative 5´ and 3´ splice sites and alternative first and last exons.

PCA was performed on alternative splicing and gene expression quantifications. A tumour-stage-independent separation between tumour and normal samples based on alternative splicing is particularly evident (Figure S1A-C) and consistent with previous studies (7, 8). Some of the events reported as significantly altered by those studies overlap those highlighted in our analysis (Figure S1B), including *RPS24* alternative exon 6, more excluded in multiple cancer types (8) and considered a potential driver of hepatocellular carcinoma (49).

Nonetheless, this strong tumour-stage-independent separation may be undermining splicing alterations that discriminate the initial stages of tumour progression, i.e. changes that contribute specifically to the separation between normal and tumour stage I samples. Therefore, PCA was performed on the alternative splicing quantification and gene expression data from the 181 tumour stage I and 112 normal breast samples (Figure 2A,B and S1D, respectively). Principal component 1, the most explanatory of data variance, separates these two groups for both PCA on alternative splicing quantification and gene expression. Several alternative splicing events contribute for that separation, including those in genes *LRRFIP2, MCM7, NUMB, SLK, SMARCC2* and *SLMAP* (Figure 2B), which also separate between all tumour stages and normal samples (Figure S1A,B and Tables S1,S2).

**Figure 2.**
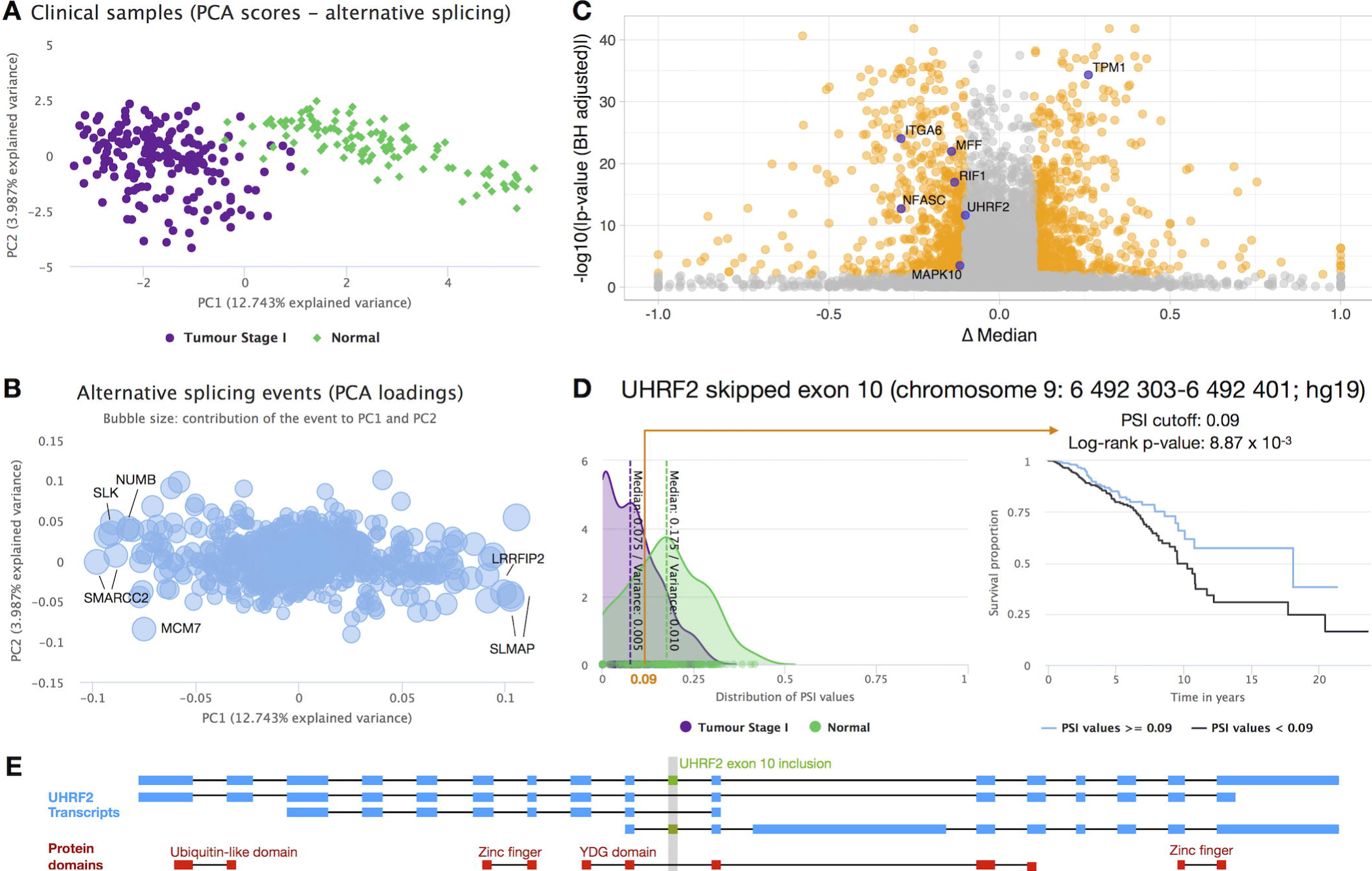
Alternative splicing analyses on tumour stage I and normal breast cancer samples from TCGA. **A,B** – PCA on PSI levels from tumour stage I and normal breast cancer samples; score (**A**) and loading (**B**) plots. The loading plot depicts the projection of splicing events on the two first principal components, with selected events labelled with their cognate gene symbol. The bubble size in panel B represents the relative contribution of each alternative splicing event to the selected principal components. **C -** Volcano plot of differential splicing analysis performed between tumour stage I and normal breast cancer samples using the Wilcoxon rank-sum test with Benjamini-Hochberg (FDR) adjustment for multiple testing. Significantly differentially spliced events (|∆ median PSI| > 0.1 and FDR ≥ 0.01) are highlighted in orange, with selected events with putative prognostic value depicted in purple. **D** – One such event is an *UHRF2* skipped exon, whose PSI distributions in tumour stage I and normal samples are depicted in the density plot (left), whereas its prognostic value is illustrated by the Kaplan-Meier survival curves (right; patients separated by a PSI cut-off of 0.09). **E -** Protein domain disrupted by *UHRF2* exon inclusion. *UHRF2* transcripts in blue, *UHRF2* exon 10 in green and UniProt domains in red. All images were retrieved from *psichomics* as is, with the exception of the gene symbol overlay in B and C, the arrow highlighting the PSI cut-off in D and panel E.

SLMAP, for instance, is a membrane protein suggested to be a mediator of phagocytic signalling of macrophages and a putative biomarker in drug-resistant cancer cells (50). *SLMAP* exon 23 encodes the protein’s tail anchor and thus its splicing, which our analyses find altered in breast stage I tumours, determines its subcellular localisation (51).

We also detected alterations in the splicing of exon 12 in *NUMB*, whose protein is crucial for cell differentiation as a key regulator of the Notch pathway. Of note, the RNA-binding protein QKI has been shown to repress *NUMB* exon 12 inclusion in lung cancer cells by competing with core splicing factor SF1 for binding to the branchpoint sequence, thereby repressing the Notch signalling pathway, which results in decreased cancer cell proliferation (52). Consistently, when analysing all TCGA breast normal and tumour samples, we show *NUMB* exon 12 inclusion is increased in cancer and negatively correlated with *QKI* expression (Spearman’s *rho* = −0.549, p-value < 0.01; Figure S2).

Complementary analyses deemed 1285 events to be differentially spliced between tumour stage I and normal samples (|∆ median PSI| > 0.1 and FDR ≤ 0.01, Benjamini-Hochberg adjustment to Wilcoxon rank-sum test; Figure 2C and Table S5) and therefore potential biomarkers for early breast cancer diagnosis. Some of the identified events (for instance, in *FBLN2* and *AP2B1)*, have already been described as oncogenic drivers following experimental validation (8).

Next, several alternative splicing events potentially associated with prognosis were identified in *UHRF2, MAPK10, RIF1, MFF, TPM1, ITGA6* and *NFASC*, based on overall survival analyses stratified by their respective optimal PSI cut-offs (labelled in Figure 2C; survival curves in Figures 2D and S3).

Detected alterations in alternative splicing may simply be a reflection of changes in gene expression levels. Therefore, to disentangle these two effects, differential expression analysis between tumour stage I and normal samples was also performed (Figure S4). Alternative splicing changes seem to be independent from alterations in the expression of cognate genes for 4 of the 7 prognosis-associated splicing events (labelled points in Figure S4).

One of such events is the alternative splicing of *UHRF2* exon 10. Cell-cycle regulator UHRF2 promotes cell proliferation and inhibits the expression of tumour suppressors in breast cancer (53). *psichomics* reveals that higher inclusion of *UHRF2* exon 10 is associated with normal samples and better prognosis (Figure 2D), and potentially disrupts UHRF2’s SRA-YDG protein domain, related to the binding affinity to epigenetic marks (Figure 2E). Hence, exon 10 inclusion may supress UHRF2’s oncogenic role in breast cancer by impairing its activity through the induction of a truncated protein or a non-coding isoform (Figure 2E). Moreover, this hypothesis is independent from gene expression changes, as *UHRF2* is not differentially expressed between tumour stage I and normal samples (|log_2_(fold-change)| < 1; Figures S4 and S5A) and there is no significant difference in survival between patient groups stratified by its expression in tumour samples (log-rank p-value = 0.279; Figure S5B).

To our knowledge, the putative prognostic value of *UHRF2* exon 10 has never been described and, together with the finding of both novel and previously validated cancer-specific alternative splicing alterations, demonstrates the potential of *psichomics* in uncovering alternative splicing-related molecular mechanisms underlying disease and physiological conditions.

### Performance benchmark

The time required to load, quantify and analyse data from different TCGA and GTEx cohorts was benchmarked. The breast cancer cohort contains the highest number of RNA-seq samples available in TCGA, thus being that for which it takes more time to load, quantify and analyse alternative splicing and gene expression data. Contrastingly, processed data from GTEx come bundled in files containing all tissues. Although only data from specified tissues are loaded, scanning though the large GTEx file still delays data loading. Tissues from GTEx were loaded in pairs for subsequent differential splicing analyses (Figure 3A).

Synthetic datasets for gene expression and exon-exon junction quantification of multiple sample sizes were generated, based on TCGA data distributions, to determine the time complexity of each step in *psichomics* as a function of the number of input samples s (Figure 3B). Assuming a constant number of genes (20000 in the benchmark) or exon-exon junctions (200000), the time taken to load data grows quadratically with s. Gene expression normalisation and differential expression are based on commonly-used, time-efficient bioinformatics tools and the times taken for each also grow quadratically with s. Alternative splicing quantification is associated with element-wise operations on matrices of dimensions s by the number of alternative splicing events and takes a runtime approximately proportional to the square of s, for a given number of alternative splicing events (around 9000 for each benchmarked run). Finally, differential splicing is based on multiple, distinct statistical analyses of alternative splicing quantification data and grows linearly with s.

## DISCUSSION

Alternative splicing is a regulated molecular mechanism involved in multiple cellular processes and its dysregulation has been associated with diverse pathologies (1-3, 5). The advent of next-generation sequencing technologies has allowed the investigation of transcriptomes of human biological samples to be expanded to alternative splicing. RNA-seq data, like those yielded by the GTEx and TCGA projects, are indeed playing crucial role in the improvement of our insights into the role of alternative splicing in both physiological and pathological contexts (2, 3, 6-8).

However, the most commonly used tools for alternative splicing analyses currently do not allow researchers to fully benefit from the wealth of pre-processed RNA-seq data made publicly available by the aforementioned projects. For instance, they lack support for estimating PSIs based on splice junction read counts. Such functionality would allow users to overcome the difficulties caused by the raw RNA-seq data from GTEx and TCGA being under controlled access and, more importantly, their processing requiring computational resources inaccessible to the majority of research labs. *psichomics* thus exploits pre-processed alternative splicing annotation and exon-exon junction read count data from TCGA and GTEx, two of the richest sources of molecular information on human tissues in physiological and pathological conditions, as well as user-owned data, allowing researchers to hasten alternative splicing quantification and subsequent analyses by avoiding the time-consuming alignment of RNA-seq data to a genome or transcriptome of reference followed by splice junction detection.

Together with support for the integration of molecular and sample-associated clinical information, the group creation functionalities featured in *psichomics* ensure full customisability of data grouping for downstream analyses. Interesting groups to compare in TCGA, for instance, may range from the simple contrast between reformed and current smokers in lung cancer to complex combinations of gender, race, age, country and other subject attributes across multiple cancers. When survival data are available, survival analyses can be performed on samples by PSI or gene expression levels, thereby assessing the putative prognostic value of a respective molecular feature.

The integrative analysis of publicly available TCGA data by *psichomics* allowed us to identify multiple exons differentially spliced between breast tumour stage I and normal samples, therefore deeming them potential diagnostic biomarkers, and to assess their putative prognostic value. The output of *psichomics* is validated by identified alternative splicing alterations that have been previously linked to the disease, including events in *RPS24, NUMB, FBLN2* and *AP2B1*. Previously understudied, yet intriguing, events were also identified, such as the skipping of SLMAP exon 23 and UHRF2 exon 10. These may provide novel insights into the early stages of breast cancer development. Indeed, it is of utmost importance to foster alternative splicing analyses of clinical samples as a crucial complement to more conventional research focused on total gene expression.

To ensure researchers with different skills can take the most out of *psichomics*, users lacking a computational background may feel more comfortable using the intuitive and more accessible graphical interface, whereas advanced users may opt for the command-line view.

Notwithstanding its merits, a current limitation of *psichomics* is the current support only for events quantified based on exon-exon junction read counts, as not all types of alternative splicing events can be profiled using splice junction reads alone. For instance, exon-intron junction, exon body and intron body quantifications are vital to confirm intron retention and alternative 5´ and 3´ UTR events over further transcriptional variations (27, 54). However, although GTEx (but not TCGA) readily provides intron and exon body read quantification for retrieval, neither TCGA nor GTEx provide exon-intron junction quantification. As input data may also be user-provided, we are developing support for the missing types of events to be included in a future update.

Further developments of *psichomics* are in progress, including alignment of raw RNA-seq data preceding alternative splicing quantification and automatic data retrieval and processing from additional public sources, such as *recount2*, an online resource containing processed data (including splice junction quantifications) for 2041 RNA-seq studies (55).

Using *psichomics*, we are able not only to identify novel exons differentially spliced between tumour stage I and normal breast samples but also to pinpoint potentially clinically relevant splicing events by embracing clinical data and evaluating their prognostic value. We expect that fellow researchers and clinicians will be able to intuitively employ *psichomics* to assist them in uncovering novel splicing-associated prognostic factors and therapeutic targets, as well as in advancing our understanding of how alternative splicing is regulated in physiological and disease contexts.

## AVAILABILITY

*psichomics* is an open-source R package publicly available in Bioconductor at https://bioconductor.org/packages/psichomics, along with graphical and command-line interface tutorials based on the presented case study.

## ACCESSION NUMBERS

Not applicable.

## SUPPLEMENTARY DATA

Supplementary Data are available at NAR online.

## ACKNOWLEDGEMENT

We thank Lina Gallego, Marie Bordone, Maria Teresa Maia, Mariana Ferreira, Ana Carolina Leote and Bernardo de Almeida for testing the program, contributing for its improvement, and all their suggestions on the manuscript. We thank Bárbara Caravela and Susana Vinga for discussions and implementation solutions on variance-based statistical tests. We also thank Ana Rita Grosso and André Falcão for valuable suggestions during the software’s development.

